# Functional Genomics Validation of PPAR gamma Signalling in PASMCs: Therapeutic Implications for Pulmonary Arterial Hypertension

**DOI:** 10.1101/2024.12.03.626672

**Authors:** Renzhi Su, Armalya Pritazahra, Shirin C.C. Saverimuttu, Lucie H Clapp, Ruth C Lovering, Jigisha A Patel

## Abstract

Gene Ontology (GO) is a tool which provides gene functional annotations, an essential resource for knowledge discovery and the analysis of biological datasets. Although considerable research has quantified the functional similarity between gene products and physiological processes, there is a need to identify which of these may contribute to pathophysiological states in humans. Previous studies have identified the role of PPARG in multiple signalling pathways, particularly those of TGFB1 and BMP2, in pulmonary artery smooth muscle cells and their relation to pulmonary arterial hypertension (PAH). To enhance the description of PPARG in the GO resource we systematically curated the proteins it interacts with and its physiological role in PASMCs. In addition, we curated the microRNAs that play a role in PAH through their regulation of PPARG expression and their downstream impact on cellular processes. This project curated experimental evidence describing 101 human miRNAs that regulate the expression of 17 PPARG signalling pathway-relevant proteins. Of these, 91 of our curated miRNAs were previously unannotated in terms of directly regulating the expression of these priority proteins. By submitting these annotations to the GO Consortium database, we have significantly expanded the breadth and depth of the GO description of PPARG-associated signalling pathways.

## Introduction

Peroxisome proliferator-activated receptor gamma (PPARG, PPARγ) belongs to a subfamily of ligand-activated nuclear receptors. Following activation by endogenous or synthetic ligands, PPARG binds to DNA-specific PPAR response elements and modulates the transcription of its target genes ^1^. Since its discovery, PPARG has been shown to be involved in regulating many physiological processes, including lipid and glucose metabolism, inflammation, cell proliferation and blood pressure ^2^. As a master regulator, PPARG appears to increase insulin sensitivity, as evident by dominant-negative mutations in the ligand-binding domain of PPARG, leading to severe insulin resistance in subjects with type 2 diabetes ^3^. In contrast, neuronal PPARG deletion in mice diminishes hyperphagic effects and energy expenditure, demonstrating an essential role contributing to weight gain in obesity ^4^. Additionally, *in vivo* studies have demonstrated liver steatosis is exacerbated in obese mice expressing high hepatic levels of PPARG ^5^. In terms of inflammation, PPARG inhibits nuclear factor kappa B (NF-κB) through transrepression, leading to the downregulation of the pro-inflammatory genes, such as interleukin (IL) 1, 6, and 12 ^2,6^. In other studies, PPARG upregulates anti-inflammatory genes, such as IL1 receptor antagonist (IL1RN), to further counter inflammation ^7^. PPARG helps regulate vascular remodelling by inhibiting the proliferation of pulmonary arterial smooth muscle cells (PASMCs), a crucial factor in pulmonary arterial hypertension (PAH) (Figure 1)^8^.

**Figure 1.**
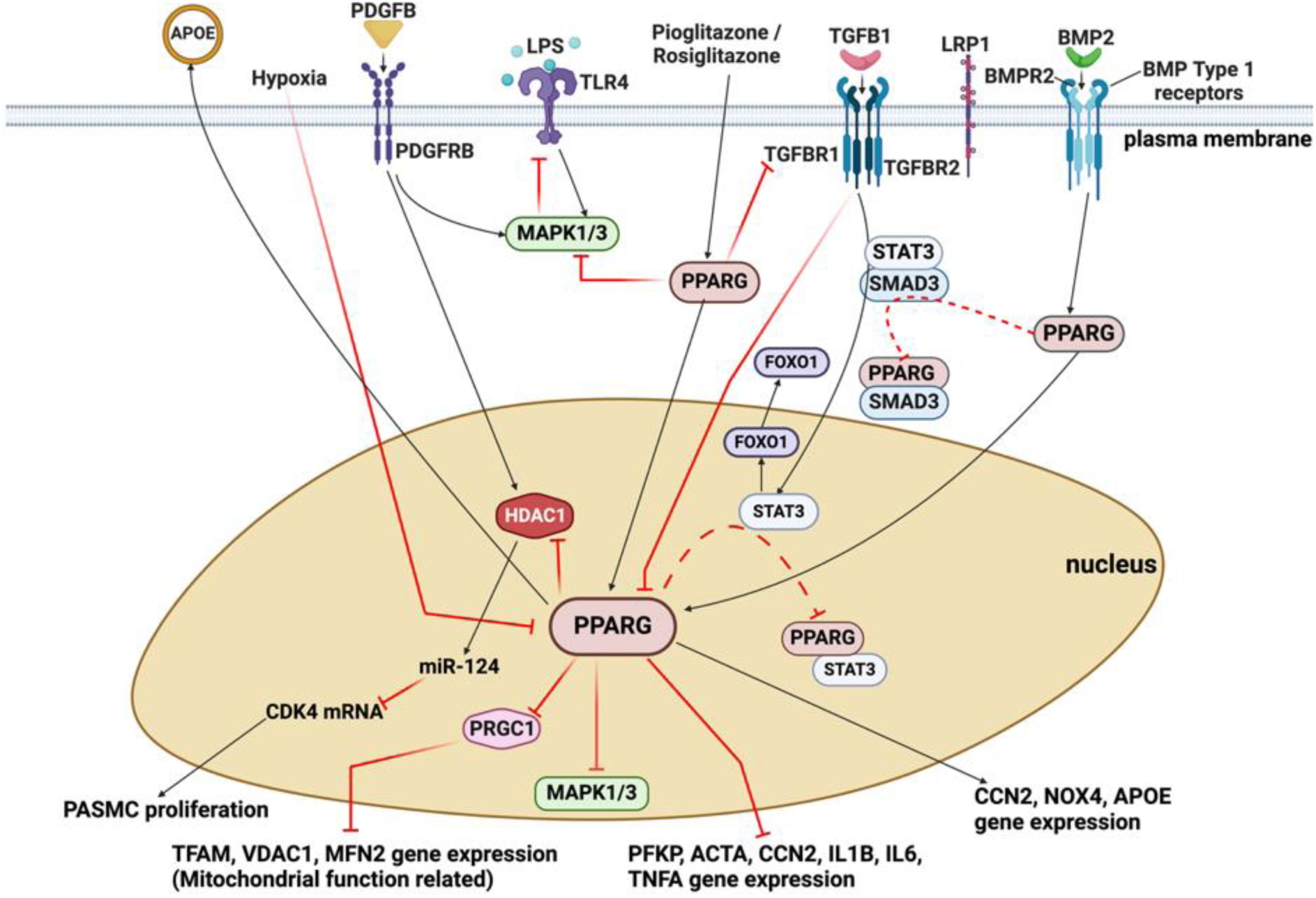
The mechanistic basis of PPARG signalling in human PASMCs. Diagrammatic representation of interactions between various signalling pathways, their downstream impact on gene regulation and other cellular processes as assessed in several publications ^8,13,15,18–21^. The focus was on the proteins associated with BMP2, TGFB1 and PPARG signalling. Black arrows illustrate activation of gene expression or positive regulation, whereas red T-shaped lines illustrate negative regulation of a biological function. The red T-shaped dotted lines indicate the physical interaction of the gene targets. This figure was generated using BioRender.com.

Synthetic PPARG ligands of the thiazolidinedione class, such as rosiglitazone and pioglitazone, are used as anti-diabetic drugs due to their potent insulin-sensitizing effects ^9^. Both rosiglitazone and pioglitazone are known to increase plasma levels of insulin-sensitizing adipokine, adiponectin ^10^. Furthermore, ligand activators of PPARG can reduce the severity of rodent models of PAH while smooth muscle-specific PPARG knock-out mice develop PAH spontaneously ^11^. However, the side effect profile of some thiazolidinediones, such as body weight gain and oedema, challenge their clinical application. Therefore, their use in the treatment of PAH warrants further investigation ^12^.

Undoubtedly multiple signalling pathways contribute to vascular remodelling in PAH (Figure 1) ^8^. As a master regulator, PPARG can crosstalk with transforming growth factor beta 1 (TGFB1, TGFβ) receptor and bone morphogenetic protein receptor type 2 (BMPR2) signalling pathways, limiting activation of the former while acting in concert with the latter to inhibit gene expression that drives inflammatory remodelling in PAH ^13^. PPARG activates phosphatases that directly inactivate [mitogen-activated protein kinase 1](https://www.genenames.org/data/gene-symbol-report/#!/hgnc_id/HGNC:6871) and 3 (MAPK1/3) and also inhibits STAT3 signalling. The key role of BMPR2 in this pathophysiology is evident, as loss-of-function mutations in BMPR2 occur in around 70% of patients with familial PAH ^14^. The dysfunction of BMPR2 could lead to a decrease in endogenous PPARG activity, resulting in the enhancement of [platelet derived growth factor subunit B](https://www.genenames.org/data/gene-symbol-report/#!/hgnc_id/HGNC:8800) (PDGFB) and MAPK signalling, and thus an increase in smooth muscle cell (SMC) proliferation, and ultimately development of PAH ^15^. Furthermore, [bone morphogenetic protein 2](https://www.genenames.org/data/gene-symbol-report/#!/hgnc_id/HGNC:1069) (BMP2) activates BMPR2 to inhibit SMC proliferation via PPARG and apoE ^15^. Transforming growth factor beta 1 (TGFB1, TGFβ) also appears to play a role in the development of PAH as TGFB1 expression is increased in PASMCs derived from PAH patients, which induces the proliferation of these cells and inflammation ^16,17^.

In addition to the protein interactions associated with PPARG signalling, microRNAs (miRNAs) also play a crucial role in regulating this pathway. By binding to the 3’ untranslated region of mRNAs, miRNAs increase mRNA degradation or inhibit translation, thus regulating gene expression ^22^. For example, miR-27a-3p binds to PPARG mRNA, leading to reduced PPARG protein levels and increased human pulmonary arterial endothelial cell proliferation ^23^. Therefore, to fully understand the development of PAH, further investigation into the systems regulating the PPARG signalling network is required. These investigations need to identify not only those miRNAs which directly target and suppress the expression of PPARG mRNA, but also the miRNAs that target other genes associated with this pathway. One way of achieving this is through *in silico* analysis using Gene Ontology (GO) data.

GO is a powerful resource that has established a standardised annotation system applying specific terminology to describe cellular processes and their relationships from the perspective of logical structure ^24,25^. In addition, a research article’s evidence of target genes and their relationship can be presented in a network to highlight crucial physiology networks ^24^. GO is used to summarise the physiological role of gene products from millions of published research articles available to search and access via databases such as PubMed and Europe PubMed Central ^26^. By condensing the knowledge of a biological entity into GO annotations, this can make biological knowledge more accessible in human and computer-readable formats. With well-structured data, the benefit to researchers is that they can extract the curated experimental evidence and plan future scientific research with organised evidence. With free access, the GO database allows all researchers to obtain the comprehensive information on target genes through public platforms, such as QuickGO and AmiGO ^24^. Additionally, these annotations are imported into the majority of tools used to analyse RNA sequencing, proteomic, transcriptomic and genomic data.

To enhance the description of human PPARG signalling in the GO resource and its physiological role in PASMCs, we systematically curated experimentally verified PPARG protein interactions. In, addition, we curated the biological roles of miRNAs that may play a role in PAH through their interaction with mRNAs encoding proteins within or associated with the PPARG signalling pathway. We limited the curation focus to two major signalling pathways (those activated by BMP2 and TGFB1), linked to PPARG and known to be dysregulated in PAH ^8,13,15,18–21,27^. In summary, our project curated experimental evidence describing 101 human miRNAs that regulate the expression of 17 PPARG signalling pathway-relevant proteins. Out of these, 91 miRNAs were previously unannotated in terms of directly regulating the expression of these priority proteins. Therefore, by submitting these annotations to the GO Consortium database, we have significantly expanded the breadth and depth of the GO description of PPARG-associated signalling pathways.

## Methods

### Curation priorities

The overall focus of this study was to identify and curate experimental evidence published before 30^th^ July 2023, describing miRNAs that regulate the expression of PPARG signalling pathway-relevant human proteins. This approach led to 17 proteins being prioritised for investigation: ACVRL1, activin A receptor like type 1; BMP2, bone morphogenetic protein 2; BMPR1A (ALK3), BMPR1B, BMPR2 (BMPR-II), bone morphogenetic protein receptors 1A, 1B and 2; PPARG (PPARγ), peroxisome proliferator activated receptor gamma; SMAD1, SMAD2, SMAD3, SMAD4, SMAD5, SMAD9 (SMAD8), SMAD family members 1-5 and 9; STAT3, signal transducer and activator of transcription 3; TGFB1, transforming growth factor beta 1 (TFGβ); TGFBR1, TGFBR2, and TGFBR3, transforming growth factor beta receptors 1-3 (Supplemental Table1).

### Identification of publications describing miRNAs regulating the expression of priority list genes

Having prioritised 17 target proteins, two miRNA text mining databases were used to identify the majority of articles for curation, namely emiRIT (extracting miRNA Information from Text) and MiRTarBase ^28,29^. These resources were searched using Human Genome Organisation (HUGO) Gene Nomenclature Committee (HGNC) approved gene symbols for the priority miRNA targets ^30^. Only articles describing the role of human miRNAs were reviewed. In addition, only those that included experimental evidence of a direct miRNA-to-mRNA interaction (usually by a luciferase assay) and confirmed that the miRNA regulated the expression of the target mRNA and/or protein were curated. In two cases, ACVRL1 and STAT3, articles were identified in PubMed using the search terms: ‘ACVRL1 + microRNA’ and ‘STAT3 + microRNA + luciferase + pulmonary’. For ACVRL1, this increased the number of articles to curate, but in the case of STAT3, the number of articles identified was reduced from over 400 to 67.

### Curation process

The research articles selected in this project were curated following established GO Consortium guidelines ^31,32^. Standard GO terms were applied to describe all experimental evidence suitable for annotation in the articles curated, including the molecular functions, biological processes and cellular locations of proteins and miRNAs. As previously described, the GO annotation extension field was used to capture contextual information ^33^. In a few cases, non-human protein GO annotations were copied to the orthologous human protein records following GO Consortium guidelines and using the evidence code “Inferred from Sequence or Structural Similarity” (ISS ^32^).

### Accessing GO annotations

Annotations contributed by this project to the GO Consortium database are attributed to the British Heart Foundation and University College London collaboration (BHF-UCL). The annotations are free to access from browsers such as QuickGO and AmiGO ^24,34^ and other bioinformatic databases, including Ensembl, miRBase, NCBI gene, RNAcentral and UniProt ^35–39^. The interaction annotations contributed by this project to the GO Consortium database can also be accessed in a format compatible for use in Cytoscape ^40^. via the Proteomics Standard Initiative Common QUery InterfaCe (PSCIQUIC) web service ^41^, PSCIQUIC provides two European Bioinformatics Institute-Gene Ontology Annotation (EBI-GOA) datasets, one with protein:protein interaction data (EBI-GOA-nonIntAct) and one with miRNA:mRNA interactions (EBI-GOA-miRNA) ^42–44^.

### Molecular interaction networks

Two different molecular interaction networks were created in Cytoscape: a network of all the interaction data submitted to the GO Consortium annotation files during this study and a network limited to the miRNAs regulating the 17 priority proteins associated with the BMP2, TGFB1 and PPARG signalling pathways. Both networks were constructed using Cytoscape version 3.10.1 ^40^.

#### Project interaction network

A network of interactions based on the annotations made by this project was created manually by using Cytoscape. First, the article PubMed IDs curated during this study were used to extract the annotated data from QuickGO. Next, the data was edited in an Excel file by following the PSI-MI TAB format (https://psicquic.github.io/MITAB28Format.html) and saved as a text file (.txt). Finally, this file of molecular interaction data was imported to Cytoscape. The network was edited to remove self-loops and duplicate edges, to change the node styles to distinguish the different entity types and to provide human-readable labels rather than UniProt and RNAcentral IDs.

#### Network of miRNAs regulating BMP2, TGFB1 and PPARG signalling pathways

This molecular interaction network was constructed in Cytoscape by seeding the network with the priority list of genes (Supplementart Table 1) and importing the interacting miRNAs from the EBI-GOA-miRNA file on 8 December 2023. As mentioned above, once the network was created, all duplicate edges and self-loops were removed, the node styles were changed to distinguish mRNAs from non-coding RNAs, and human-readable labels were provided. As previously described, the GO term enrichment analysis was undertaken using the Cytoscape plugins GOlorize and BiNGO ^42–44^. The settings applied were overrepresentation, Hypergeometric test, Benjamini & Hochberg False Discovery Rate (FDR) correction, and significance level ≤0.05. The two functional enrichments were undertaken with the human GO association files downloaded on 17 December 2019 and 30 October 2023 and the obo ontology file downloaded on 24 January 2024. For each year, the protein and RNA annotation files were combined, and all available annotations were used as the reference set. Enriched GO terms with two or less associated gene products were removed from both analyses. Finally, a selection of GO terms was overlaid onto the interaction network.

**Table 1.** Selection of GO terms enriched in the functional analysis of the network of miRNAs regulating BMP2, TGFB1 and PPARG signalling. Two analyses were conducted to compare annotation data available in 2019 with the 2023 data. Bold text emphasises an enriched GO term that has at least 3 times more network gene products associated with it in 2023 than in 2019. *Enriched GO terms that are overlaid onto the network in Figure 6. The number of human gene products in the genome associated with a biological process GO term in 2019 and 2023 was 30629 and 37617, respectively. (x indicates the number of gene products in the network associated with the GO term; n indicates the number of gene products in the genome associated with a GO term).

|  | 2023 |  |  |  |  |  | 2019 |  |  |  |  |
| --- | --- | --- | --- | --- | --- | --- | --- | --- | --- | --- | --- |
| GO term name | corrected p-value | x | n | number of mRNAs | number of miRNAs |  | corrected p-value | x | n | number of mRNAs | number of miRNAs |
| iRNA-mediated post-transcriptional gene silencing | 1.63E-72 | 107 | 5097 | 0 | 107 |  | 2.38E-47 | 75 | 3866 | 1 | 75 |
| **miRNA-mediated gene silencing by mRNA destabilization** | **0.00E+00** | **73** | **107** | **0** | **73** |  | **#N/A** | **#N/A** | **#N/A** | **0** | **0** |
| regulation of cell adhesion | 2.28E-11 | 20 | 807 | 5 | 15 |  | 4.09E-08 | 15 | 714 | 5 | 10 |
| **regulation of cell adhesion molecule production** | **4.89E-10** | **6** | **21** | **0** | **6** |  | **#N/A** | **#N/A** | **#N/A** | **0** | **0** |
| **regulation of extracellular matrix assembly** | **2.44E-31** | **16** | **30** | **6** | **10** |  | **6.11E-05** | **3** | **15** | **2** | **1** |
| *regulation of cell migration | 5.19E-62 | 63 | 1061 | 11 | 52 |  | 6.65E-43 | 47 | 973 | 10 | 37 |
| regulation of blood vessel endothelial cell migration | 1.37E-42 | 30 | 165 | 4 | 26 |  | 1.02E-32 | 24 | 157 | 3 | 21 |
| regulation of vascular associated smooth muscle cell migration | 2.06E-20 | 13 | 51 | 0 | 13 |  | 2.03E-15 | 10 | 44 | 0 | 10 |
| *regulation of endothelial cell proliferation | 4.84E-37 | 28 | 189 | 4 | 24 |  | 1.35E-38 | 28 | 179 | 6 | 22 |
| *regulation of smooth muscle cell proliferation | 3.39E-27 | 22 | 177 | 5 | 17 |  | 3.50E-23 | 19 | 169 | 3 | 16 |
| *apoptotic process | 1.60E-02 | 9 | 1035 | 6 | 3 |  | 3.97E-02 | 7 | 903 | 5 | 2 |
| *regulation of apoptotic process | 1.11E-23 | 40 | 1557 | 8 | 32 |  | 2.72E-16 | 31 | 1541 | 5 | 26 |
| cellular response to hypoxia | 1.51E-11 | 11 | 141 | 2 | 9 |  | 5.24E-08 | 8 | 128 | 0 | 8 |
| regulation of cytokine production | 2.60E-54 | 55 | 884 | 5 | 50 |  | 6.46E-24 | 30 | 750 | 4 | 26 |
| *transforming growth factor beta receptor superfamily signaling pathway | 5.25E-24 | 20 | 174 | 17 | 3 |  | 1.38E-21 | 18 | 167 | 15 | 3 |
| **\*regulation of transforming growth factor beta receptor signaling pathway** | **1.92E-63** | **41** | **179** | **7** | **34** |  | **3.94E-11** | **10** | **117** | **7** | **3** |
| *BMP signaling pathway | 3.97E-16 | 12 | 77 | 11 | 1 |  | 8.78E-16 | 12 | 86 | 11 | 1 |
| *regulation of BMP signaling pathway | 2.58E-25 | 19 | 123 | 3 | 16 |  | 1.07E-07 | 7 | 91 | 4 | 3 |
| SMAD protein signal transduction | 7.12E-10 | 7 | 42 | 7 | 0 |  | 1.16E-10 | 8 | 58 | 8 | 0 |
| **regulation of SMAD protein signal transduction** | **1.14E-49** | **28** | **75** | **10** | **18** |  | **5.92E-06** | **4** | **25** | **4** | **0** |

## RESULTS

### Annotation Summary

This project aimed to curate miRNAs that may have a role in PPARG signalling pathways in human PASMCs. The review of seven key references led the study to focus on the BMP2, TGFB1 and PPARG signalling pathways (Figure 1 ^8,13,15,18–21^). Following an expanded review of the literature, 17 proteins selected were prioritised for further investigation: BMP2, BMPR1A, BMPR1B, BMPR2, PPARG, SMAD1, SMAD2, SMAD3, SMAD4, SMAD5, SMAD9, STAT3, TGFB1, TGFBR1, TGFBR2, and TGFBR3 (Figure 2). A total of 133 articles, which described experimental confirmation of miRNA interactions with the mRNA encoding at least one of these 17 proteins, were fully curated ^42^. Following the GO Consortium guidelines ^31,32^, 966 human gene annotations were generated utilising 169 different GO terms. These annotations captured the biological roles of 108 miRNAs, of which 101 miRNAs were found to regulate the expression of the 17 priority proteins directly. In total, there are now 154 miRNA:mRNA target interactions available in the GO Consortium annotation and PSCIQUIC interaction files that capture the role of 107 miRNAs that have the potential to regulate BMP2, TGFB1 and PPARG signalling (Figure 3). This project has captured the majority of this data. However, previous work had captured the role of 16 miRNAs directly targeting one or more of the 17 priority protein mRNAs (Figure 3). Thus, this project has substantially increased the number of experimentally verified miRNA:mRNA interactions available in public bioinformatic resources associated with the BMP2, TGFB1 and PPARG signalling. Furthermore, as this project undertook a full article curation approach, many additional interactions were captured, providing breadth as well as depth to the GO Consortium resource. In total, data describing 108 miRNAs regulating the expression of 100 proteins were captured, as well as 11 interactions between miRNAs and long non-coding RNAs (lncRNAs) and 12 interactions between mRNAs and lncRNAs (Figure 4).

**Figure 2.**
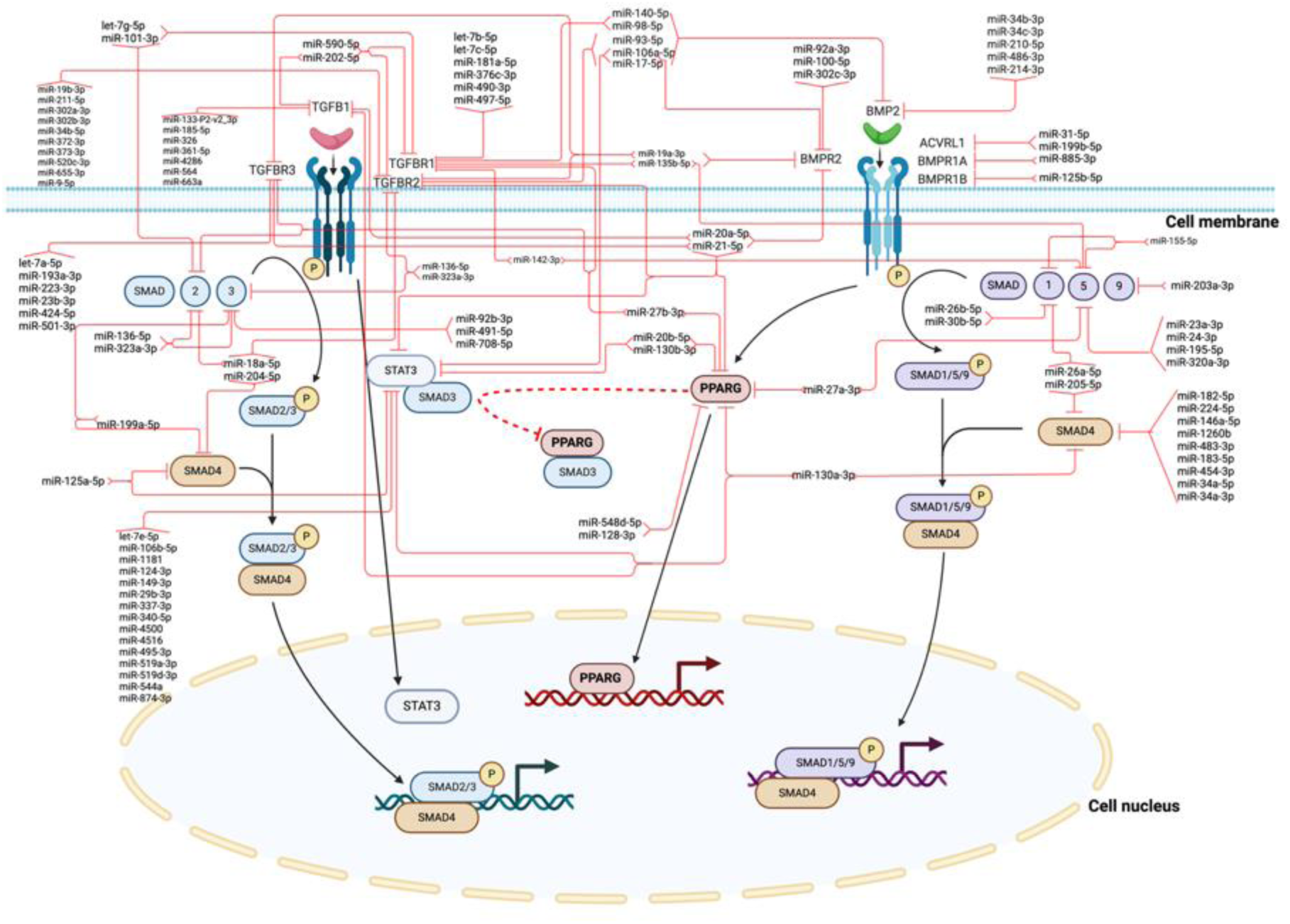
Regulation of BMP2, TGFB1 and PPARG signalling by miRNAs. This schematic represents signalling pathways based on data extracted from 133 articles describing the impact of human miRNA:mRNA target interactions regulating signalling pathways of BMP2, TGFB1 and PPARG; 154 were captured through the GO curation of 133 articles. Only direct miRNA:mRNA target interactions with experimental evidence confirming a change in expression of the target mRNA and/or protein were included. For simplicity, the direct miRNA targets are only indicated once in the pathway, even if the miRNA target could be represented more than once in the schematic. Black arrows illustrate the activation of gene expression or positive regulation, whereas red T-shaped lines illustrate negative regulation of a biological function. The red T-shaped dotted lines indicate the physical interaction of the gene targets. Pink T-shaped lines illustrate the inhibition of target gene expression by miRNAs. This figure was generated using BioRender.com.

**Figure 3.**
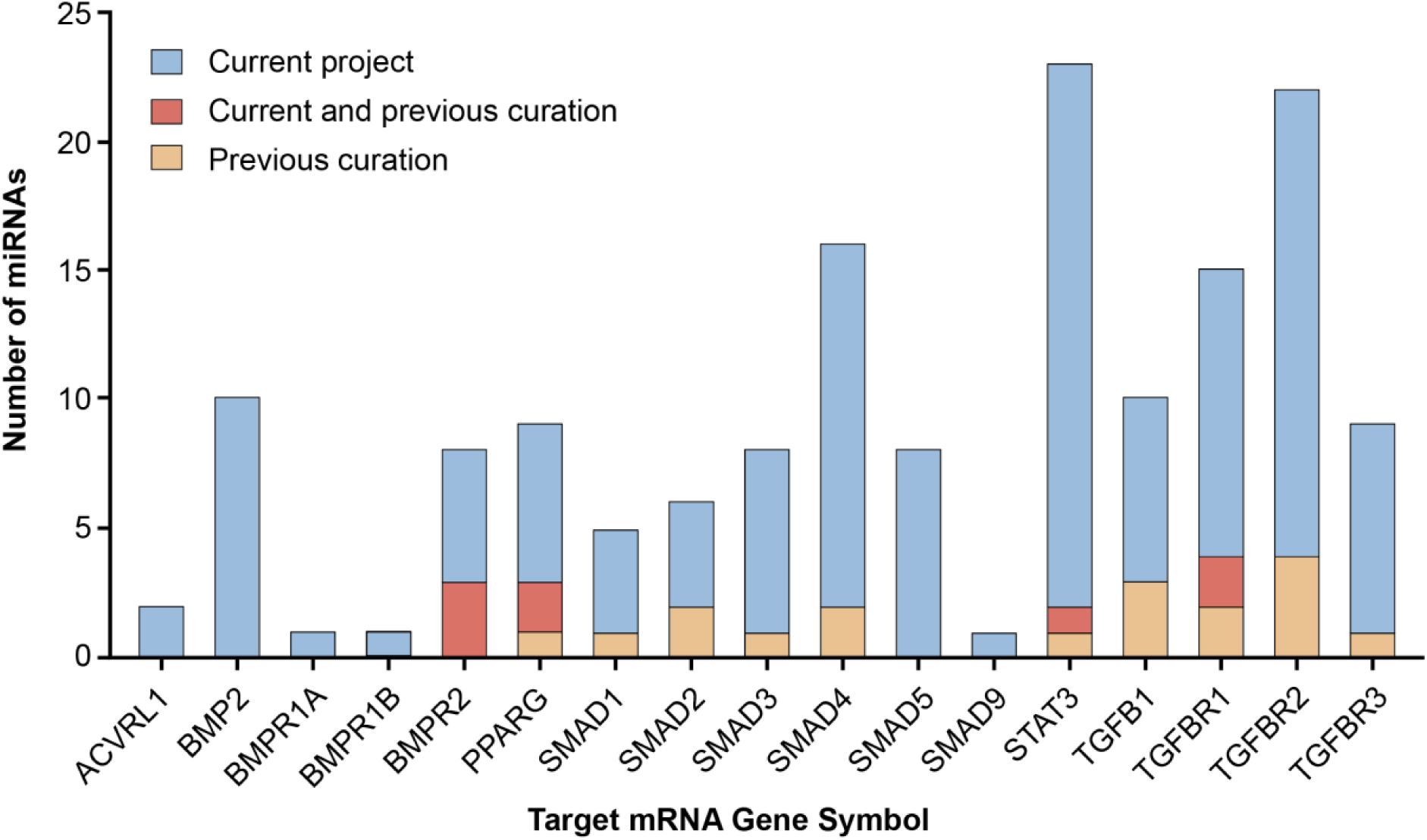
Number of miRNAs regulating the expression of 17 priority proteins involved in BMP2, TGFB1 and PPARG signalling. This current project provides 128 new miRNA:mRNA target interaction annotations (blue). 26 miRNA:mRNA target interactions were created by previous work, of which 8 were also captured by the current project (red) and 18 were exclusively captured by previous work (orange).

**Figure 4.**
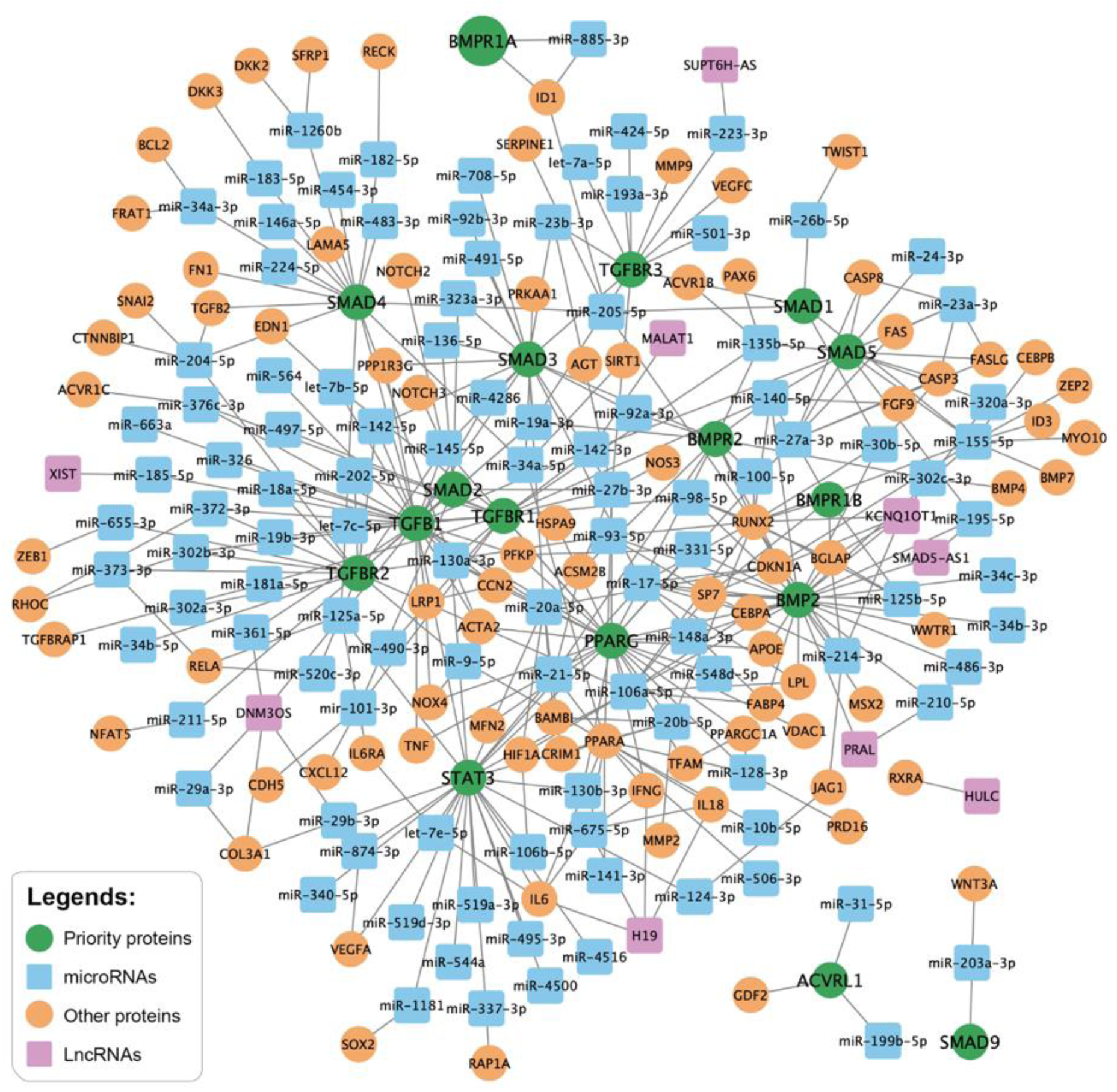
Cytoscape network analysis of interactions captured by the current project. The figure demonstrates a network of new miRNA:mRNA target interactions captured by the current project following the curation of 133 articles. This project also captured new long non-coding RNA (lncRNA) interactions with miRNAs and mRNAs. (Green round nodes: mRNAs encoding priority proteins; Blue square nodes: miRNAs; Orange round nodes: mRNA encoding non-priority proteins; pink square nodes: lncRNAs; Black edges: interactions between two nodes).

### Impact of this project

A miRNA:mRNA molecular interaction network was created to examine the impact of the annotations created by this project. The network was seeded with the 17 priority genes and the molecular interactions downloaded from the EBI-GOA-miRNA interaction dataset on 8 December 2023 from PSICQUIC. No miRNA:mRNA interactions were available from PSICQUIC in the IMEx and IntAct interaction files for the priority list on this date. The EBI-GOA-miRNA dataset of experimentally supported miRNA:mRNA target interactions is regularly exported from the GO Consortium association file. After removing duplicate edges, the resulting network included all 107 miRNAs (nodes) and contained 154 interactions (edges). Two GO analyses of this network were then conducted using annotation data released in 2019 and 2023, with both using the January 2024 ontology. The 2019 analysis led to the overrepresentation of 906 GO terms, whereas 1037 were overrepresented in the 2023 analysis.

The impact of this project on the 107 miRNAs and the 17 priority proteins GO annotations was evident when the functional analyses conducted with data retrieved from December 2019 were compared to those performed using the updated GO annotation data from October 2023. In 2019, 30 of the network genes had no GO biological process annotations, while in 2023, all 124 genes were associated with a biological process GO terms. Many of the enriched GO terms present in both analyses now have an increased number of our network gene products associated with them: 17 and 42 GO terms now have 3 and 2 times (respectively) more gene products associated with them than in 2019 (Supplementary Table 2). For example, although the GO term ‘regulation of SMAD protein signal transduction’ is significantly enriched in both 2019 and 2023 datasets, the number of genes associated with this term has increased from 25 to 75 over the past four years (Table 1). This increase has impacted our network analysis; in 2019, only 4 genes were associated with this term, whereas now there are 28. Of these 28 genes, 18 encode miRNAs, and 10 encode proteins. In addition, while 55 GO terms enriched in 2019 are not enriched in 2023, 186 GO terms that were not overrepresented in 2019 are now enriched, even though all these terms existed in 2019. Previous studies often highlight changes over time in the enriched GO terms that have been identified following a functional analysis ^45^. Similar to these studies several factors are likely to be responsible for the differences between our 2023 and 2019 analyses, including rearrangements to the ontology itself ^24^, re-annotation of existing GO annotations ^46^ or the addition of new GO annotations ^42^. The current project has focused on the BMP2, TGFB1 and PPARG signalling pathways in human PASMCs and curation of miRNAs. Consequently, for the GO terms relevant to these pathways, it is the increased number of annotations associated with these terms that is responsible for the identified changes in the number of genes associated with them.

### GO analysis of BMP2, TGFB1 and PPARG signalling pathways and associated miRNAs

The 2023 functional analysis of the miRNA, BMP2, TGFB1 and PPARG signalling pathways network enriched over 1,000 GO terms. To help interpret this data, 11 biologicals ‘general area groups’ were created, namely: cell death, cell development, cell migration, cell proliferation, gene expression, metabolic process, organelle/cellular, structure organisation, response to stimulus, signalling pathway, tissue or organ development, transport/localisation/homeostasis. In addition, a ‘high level term’ group was created for GO terms that were considered too broad to be assigned to one of the 11 biological groupings. These 12 general groups were then subdivided into 184 subgroups (Supplementary Table 3). Each enriched GO term was assigned to one of the subgroup terms to simplify the interpretation (Supplementary Table 2). This approach confirmed that 20% of the enriched GO terms fall within the tissue or organ development grouping while 11% are assigned to the signalling pathway group. Additionally, 26% of the GO terms are associated with a cell function relevant group and 11% with the response to stimulus grouping (Figure 5).

**Figure 5.**
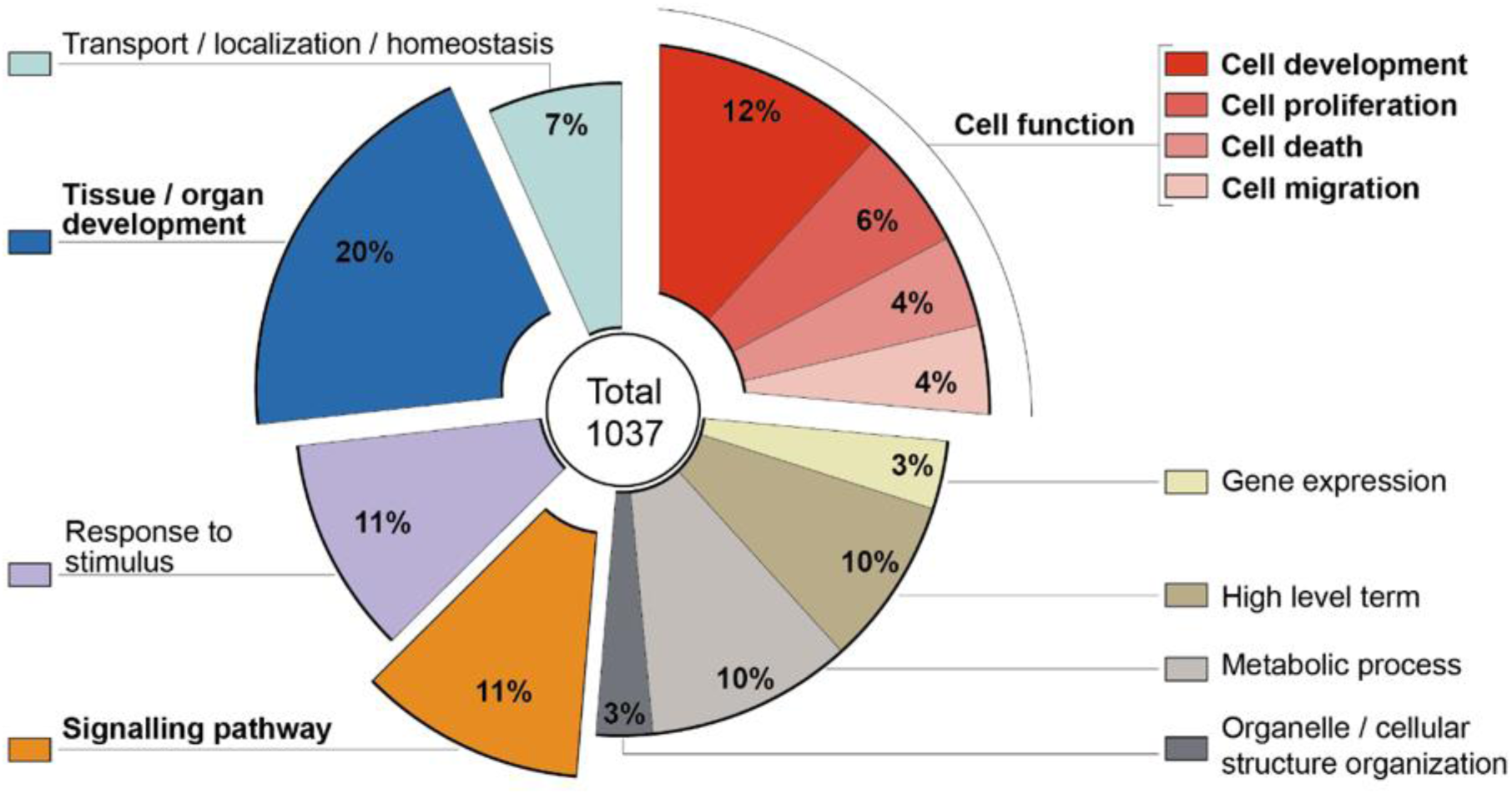
Distribution of enriched biological process GO terms within the general grouping terms. The 2023 enrichment analysis identified 1037 enriched GO terms. To simplify the interpretation of this data, terms have been classified into 12 general grouping terms (Supplementary Table 2 & 3).

### Cytoscape analysis of miRNA-target interaction network

There are a limited number of functional analysis tools that integrate miRNA GO annotations. Thus, in order to investigate the GO terms pertinent to the physiological PPARG signalling in human PASMCs and it’s regulation by miRNAs, a Cytoscape analysis was undertaken to create a network of miRNA:mRNA interactions ^40^. All miRNA:target interactions curated during this project were included in the analysis. Thus, the network was seeded with the 17 priority protein UniProt IDs. The miRNA:target network was then imported from the EBI-GOA-miRNA file, which contains experimentally validated miRNA:target data from the GO annotations files in a Cytoscape compatible format. The resulting network included 154 miRNAs:target interactions, representing 107 miRNAs and 17 protein mRNAs (Figure 6). All of these direct miRNA:target interactions are supported by experimental evidence which confirms that they regulate the expression of the target gene. The GO annotation files include UniProt IDs to denote the encoding mRNAs, rather than employing the ’mRNA identifiers’ supplied by Ensembl or NCBI. Utilising UniProt IDs in the GO Consortium annotations for human mRNAs facilitates a more accessible GO enrichment analysis of Cytoscape networks. A GO enrichment analysis was performed, and a selection of GO terms were overlaid onto the interaction network (Figure 6). The number of gene products associated with the overlaid GO terms are available in Table 1. The selected GO terms highlight the major physiological role of PPARG in PASMCs and these GO terms are also highly linked to the pathophysiological role in PAH. The GO term “miRNA-mediated post-transcriptional gene silencing” is associated with all miRNAs in this network. However, of the 107 miRNAs, 75 were associated with multiple GO terms such as "regulation of smooth muscle cell proliferation" and "regulation of apoptotic process".

**Figure 6.**
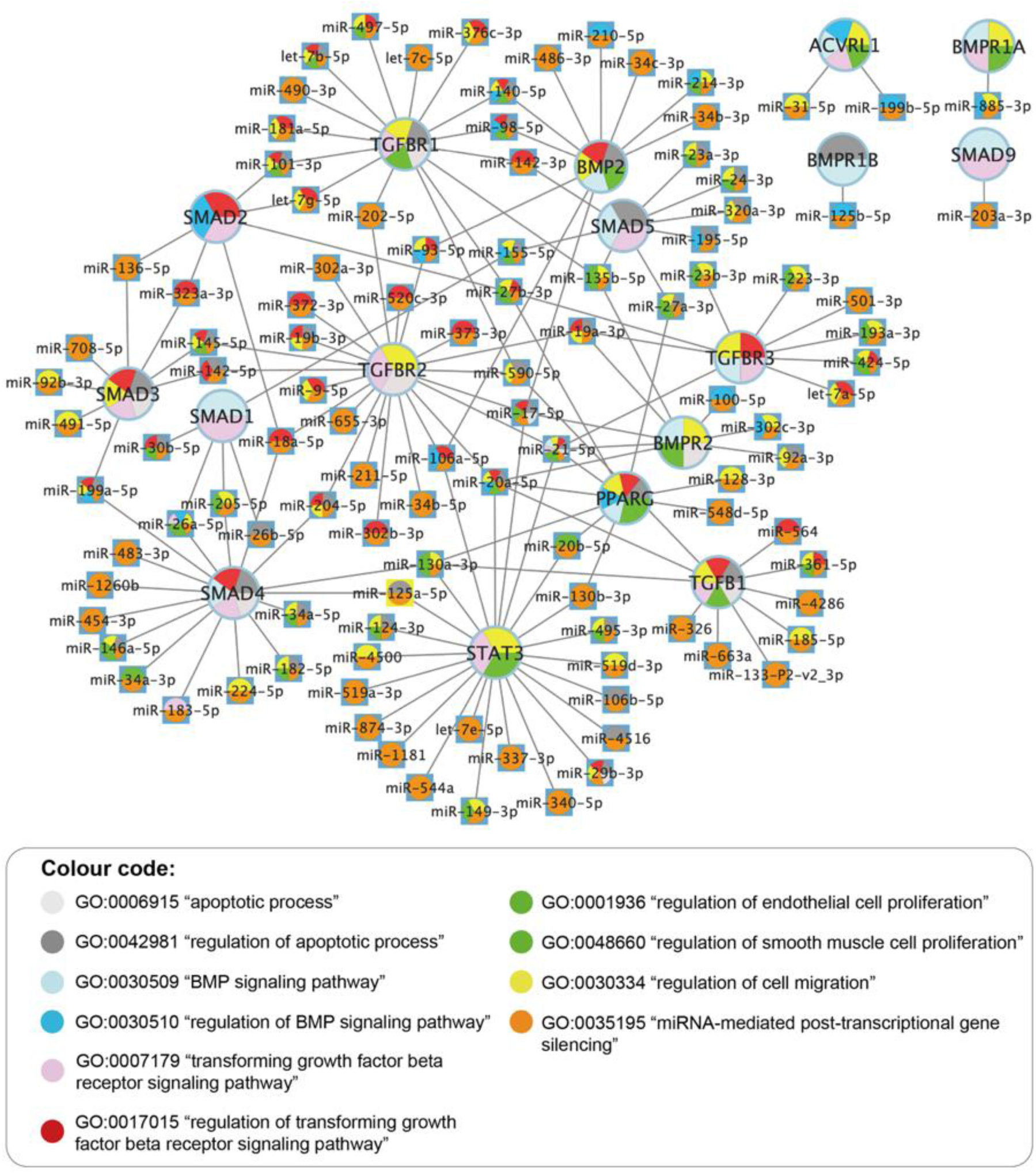
Network of miRNAs regulating BMP2, TGFB1 and PPARG signalling pathway proteins. A Cytoscape network capturing the miRNA:mRNA target interactions that are predicted to impact BMP2, TGFB1 and PPARG signalling. The network has a total of 107 miRNAs and their 17 mRNA targets encoding the priority proteins, these interactions have all been experimentally validated. Overlaid on the network are a selection of enriched GO terms (Table 1). (Round nodes: mRNAs, square nodes: miRNAs, node colours indicate overlaid enriched GO terms and edges represent interactions).

## Discussion

This investigation highlights the key physiological processes regulated by PPARG-centred signalling networks through GO. Complex and systematic impact on physiological processes was identified by integrating GO enrichment analysis into signalling networks. Previous studies in PAH have demonstrated that when one or more core targets in the network such as PPARG or BMP2, are manipulated, this can cause systemic damage leading to a pathophysiological state ^47^. To inhibit the proinflammatory remodelling process in PAH, PPARG serves to both suppress TGFB1 signalling and activate BMP2 signalling ^13^. The impact of this co-regulation is demonstrated through our GO analysis. The selection of enriched GO terms associated with the physiological role of the BMP2, TGFB1 and PPARG network are similar, but dysregulated and therefore associated with pathophysiology in PAH (Figure 6). For example, the initiation of vascular remodelling in PAH requires changes to cell adhesion, migration and proliferation, and GO terms describing these processes, "regulation of cell adhesion", "regulation of cell migration" and "regulation of cell proliferation" are all enriched in the network of miRNAs regulating the BMP2, TGFB1 and PPARG signalling pathways (Table 1). Almost half of the miRNAs in this network now associated with the GO term "regulation of cytokine production", up from a quarter in the 2019 analysis. Inflammation being a critical feature in PAH, the role of miRNAs in regulating cytokine production is an important downstream process to have been captured. Furthermore, experimental evidence suggests that decreased PPARG expression resulted in mitochondrial function deficits in human PASMCs ^8^. In addition to the role of mitochondria in cellular respiration, mitochondria also undertake the biosynthesis of reactive oxygen species, nitric oxide, fatty acids and glycoproteins. Notably, the potential impact of the BMP2, TGFB1 and PPARG signalling pathways on mitochondrial function is highlighted by the enrichment of GO terms associated with this network on known mitochondrial processes, including "regulation of reactive oxygen species metabolic process", "regulation of nitric oxide metabolic process", "regulation of fatty acid metabolic process" and "regulation of glycoprotein biosynthetic process" (Supplementary Table 3). However, further research is required to comprehensively understand how PPARG might affect PASMC metabolism and thereby promote the development of PAH.

Understanding the pathways and downstream processes that individual proteins contribute to can help predict the impact of gene variants on disease and its progression. In addition, where there is a lack of experimental evidence for all contributors to specific pathways, the role of some participants can be inferred by their interaction with entities with a known role ^48^. For example, the idea of guilt by association would imply that although there is no experimental evidence confirming that all miRNAs described in Figure 2 affect BMP2, TGFB1 and PPARG signalling, many of these miRNAs will impact these pathways. Thus, the curation of the BMP2, TGFB1 and PPARG network by this project provides the groundwork for improving the understanding of PAH pathophysiology and consequently informs the design of new therapeutic targets. Moreover, here we have captured the role of over a hundred miRNAs in the regulation of key components of this PAH relevant network. This network also confirms that many miRNAs co-regulate the expression of multiple genes in these pathways. For example, 30 miRNAs have multiple targets, in this network, with 21 functioning solely in either BMP2 or TGFB1 pathways. This provides an opportunity to investigate these miRNAs as therapeutic targets or drugs. However, 9 miRNAs (hsa-miR-20a-5p, hsa-miR-21-5p, hsa-miR-17-5p, hsa-miR-19a-3p, hsa-miR-106a-5p, hsa-miR-135b-5p, hsa-miR-93-5p, hsa-miR-98-5p and hsa-miR-140-5p) can target genes in both BMP2 and TGFB1 pathways (Supplementary Table 1). Thus, therapeutic use of these 9 miRNAs may lead to concurrent suppression of both the BMP2 and the TGFB1 pathways in the signalling network, preventing the desired counter-regulation response in the disease to which it might be targeted against. Furthermore, future PAH research can now consider the role of these miRNAs in disease progression and/or as diagnosis/prognosis biomarkers.

This study focused on the role of PPARG signalling in various human cell types, and contributed 52% of the total PPARG GO annotations. Our annotation endeavour has confirmed that, although GO offers insights into the overarching functions and cellular localisation of numerous proteins, critical processes pivotal for standard human development and cellular functions remain inadequately represented ^49,50^.

In terms of miRNA curation, this work has significantly increased the number of miRNAs regulating the PPARG-centred signalling network from 18 to 101. Understanding miRNA interactions with mRNAs is critical for unravelling disease mechanisms, where overexpressing or silencing microRNAs represents a novel gene regulation approach, particularly in developing therapeutic strategies for diseases associated with aberrant miRNA expression or their target genes ^51^. However, most miRNAs regulate multiple genes, posing the risk of off-target effects when directing therapies against specific miRNAs ^52,53^. There are many miRNA resources which provide predicted miRNA:mRNA interaction data which are not experimentally verified. While the GO Consortium has successfully organised complex experimental protein data, its curation of experimental microRNA data remains resource-limited ^54,55^. The GO Consortium has developed very specific guidelines to ensure consistent annotation of miRNAs ^31^ and the collaborative efforts between IntAct and the GO Consortium contribute validated valuable miRNA:mRNA interaction data, significantly enhancing analysis of miRNA function ^31,54^. In addition, the role of miRNAs in downstream processes is also available through GO browsers, such as QuickGO, allowing users to obtain comprehensive information ^32,34^. However, we at UCL are the only group prioritising human miRNA GO annotation and with limited resources, considerable published experimental miRNA data remains to be captured.

Utilising GO annotation within Next-Generation Sequencing (NGS) and multi-omics investigations represents a robust methodology for discerning the functional implications inherent in expansive genomic datasets. GO furnishes a standardised vocabulary for characterising genes and gene products through descriptors about their biological processes, cellular components and molecular functions. Consequently, it provides an invaluable framework for subsequent analyses concerning identifying differentially expressed genes within the context of large-scale genomic datasets. As NGS capabilities continue to advance, the significance of GO will become increasingly evident, given the pivotal role played by the comprehensiveness of database coverage and the precision of curated data in underpinning large-scale analytical endeavours. The standardised vocabulary can link genes to different databases by providing a common framework for understanding gene functions and interactions. The GO organises existing experimental data and thus enables researchers to identify enriched biological themes among the genes identified in future studies ^24^.

One might argue that a potential limitation regarding this study arises from research indicating that focused annotation projects may introduce biases in the results obtained from enrichment analyses of transcriptomic or proteomic datasets ^24^. However, this project can significantly strengthen our confidence when performing experiments such as bulk RNA sequencing and proteomics to study the impact of PPARG-related signalling in smooth muscle in PAH patient samples. Furthermore, our analysis has shown that our full article curation provides a broad range of annotations which are integrated into the existing very large corpus of GO annotation data. In addition, our analysis highlights the importance of physiological process annotation before undertaking pathophysiological studies.

Investigators should be cognizant that identical proteins often exhibit analogous functions across various developmental and signalling pathways. In this respect, there is evidence that activating the TGFB1 signalling pathway plays a protective role in pulmonary endothelium in PAH, helping maintain normal function and structure ^56^. In contrast, in PASMCs TGFB1 is pro-proliferative and, therefore, contributes to the pathological changes associated with PAH ^57^. For example, endothelial apoptosis can induce PASMC growth via TGFB1, which may lead to the remodelling of pulmonary blood vessels and the development of PAH ^58,59^. Thus, more work still needs to be done to better understand the physiological role of the signalling network this project focused on. For example, to expand the curation to other cellular components of the pulmonary artery, such as endothelial cells and fibroblasts, so that the role of PPARG in the pulmonary artery can be fully described.

## Acknowledgements

We are extremely grateful to all members of the GO Consortium who contribute to annotation and ontology development on an ongoing basis.

## Supplementary Table legends

**Supplementary Table 1. List of miRNAs targeting mRNAs encoding priority proteins.** The majority of annotations were provided by the current project, with annotations created either previously or created by the current project; previous work is indicated in the curation column.

**Supplementary Table 2. 2023 v 2019 GO analysis of Network.** Two functional enrichments of the full network of miRNAs regulating the BMP2, TGFB1 and PPARG signalling pathways were undertaken. The 2023 analysis included the human GO association files downloaded on 30 October 2023; the 2019 analysis included the human GO association files downloaded on 17 December 2019. Both analyses used the same ontology file, downloaded on 24 January 2024. General and specific grouping terms were manually applied (see Supplementary Table 3 for list of terms) to group the unique GO ID and GO term names into larger biological domains; x indicates the number of gene products in the network associated with the GO term; n indicates the number of gene products in the genome associated with the GO term; X is the number of gene products in the network associated with a biological process GO term; N indicates the number of human gene products in the genome associated with a biological process GO term.

**Supplementary Table 3. Grouping terms for enriched GO terms.** The enriched GO terms were grouped into 12 ‘general groups’: adhesion, cell death, cell development, cell migration, cell proliferation, complex clearance/assembly, gene expression, high level term, metabolic process, organelle/cellular, structure organization, regulation of catalytic activity, response to stimulus, signalling pathway, system process, tissue or organ development, transport/localization/homeostasis. These general groups were further subdivided into a total of 184 ‘specific groupings.

